# Morphological integrity of insulin amyloid-like aggregates depends on preparation methods and post-production treatments

**DOI:** 10.1101/2022.06.27.497716

**Authors:** Camilla Thorlaksen, Adriana-Maria Stanciu, Martin Busch Neergaard, Nikos Hatzakis, Vito Foderà, Minna Groenning

## Abstract

Protein aggregates are often varying extensively in their morphological characteristics, which may lead to various biological outcomes related to e.g., immunogenicity risk. However, isolation of aggregates with a specific morphology within an ensemble is often challenging. To gain vital knowledge on the effects of aggregate characteristics, samples containing a single morphology must be produced by direct control of the aggregation process. Moreover, the formed aggregates need to be in a solvent suitable for biological assays, while keeping their morphology intact. Here we evaluated the dependence of morphology and integrity of amyloid-like fibrils and spherulites on preparation conditions and post-treatment methods. Samples containing either amyloid-like fibrils or spherulites produced from human insulin in acetic acid solutions are dependent on the presence of salt (NaCl). Moreover, mechanical shaking (600 rpm) inhibits spherulite formation, while only affecting the length of the formed fibrils compared to quiescent conditions. Besides shaking, the initial protein concentration in the formulation was found to control fibril length. Surprisingly, exchanging the solvent used for aggregate formation to a physiologically relevant buffer, had a striking effect on the morphological integrity of the fibril and spherulite samples. Especially the secondary structure of one of our spherulite samples presented dramatic changes of the aggregated β-sheet content after solvent exchange, emphasizing the importance of the aggregate stability. These results and considerations have profound implications on the data interpretation and should be implemented in the workflow for both fundamental characterization of aggregates as well as assays for evaluation of their corresponding biological effects.

## Introduction

A challenge when working with biotherapeutic proteins is their inherent ability to aggregate. Protein aggregates in drug products are unwanted due to both their impact on efficacy and their potential immunogenicity risk [1-3]. This has led to an increased interest over the years of understanding the processes driving aggregation [4-7] and their resulting biological implications [8-10]. However, aggregated protein samples often consist of heterogeneous ensembles with varying characteristics, possibly having different biological effects [11]. Working with samples, which contains morphologically different aggregates complicates obtaining unequivocal conclusions from biological assays. Isolating a single morphology from the generated ensemble can be problematic, thus the aggregation process needs to be highly controlled from the early phases. Moreover, several aggregates formed *in vitro* are generated in formulations, which are not suitable for certain characterization techniques or are not readily translatable to biological assays. Solvent exchange is thus required to allow for characterization or application in biological assays and it is often achieved via centrifugation- and dialysis-based methods [12-14]. Morphological integrity before and after solvent exchange is assumed intact, yet rarely investigated. However, drastic changes in pH and salt concentration of the solvent surrounding the generated aggregates could potentially lead to destabilization of the structure.

Insulin is a biotherapeutic protein, often utilized for studying aggregation processes [6, 15, 16], due to its ability to form amyloid-like fibrils and spherulites at acidic pH and elevated temperatures [17, 18]. Amyloid-like fibrils are elongated aggregates, with a section diameter of a few nanometres and length of several micro metres [19-21]. Despite their polymorphism depending on the external conditions during formation, their internal cross β-sheet structure seems to be common for all fibrils [22]. Spherulites are spherical aggregates with a diameter spanning from a few micrometres up to several millimetres. They consist of fibril-like material radially growing from an irregular core of protein assembly [23, 24]. Due to their internal symmetry, they typically exhibit a symmetric Maltese Cross pattern when observed under cross-polarized light [24, 25]. Similar aggregation conditions often lead to a co-existence of fibrils and spherulites within the same sample [4, 7, 18, 24, 26] and the two aggregate types are difficult to separate once formed.

In this study, we present how to produce samples containing either amyloid-like fibrils or spherulites from human insulin in acetic acid solutions, by addressing the effect of salt concentration (NaCl) and mechanical shaking on the resulting aggregate morphology. Moreover, we show how the fibril length and spherulite secondary structure can be readily tuned, by adjusting protein concentration and solvent polarity, to create samples, which can be utilized for investigating the biological effect of specific aggregate characteristics on e.g., immunogenicity risk. The sample morphology was characterized by a plethora of biophysical techniques, including transmission electron microscopy (TEM), micro-flow imaging (MFI), confocal microscopy, scanning electron microscopy (SEM) and Fourier-transform infrared (FTIR) microscopy. To make the samples translatable to any biological assay, the solvents in all prepared samples were exchanged by two common procedures, either centrifugation or dialysis, into phosphate buffer saline (PBS). We found that the morphological integrity of both amyloid-like fibrils and spherulites had been affected by the procedures, highlighting the importance of evaluating aggregate characteristics both before and after solvent exchange to ensure that correct conclusions are drawn from the biological tests.

## Materials and Methods

### Materials

Acetic acid glacial (100 %, Merck KGaA, Darmstadt, Germany), ethanol absolute (≥ 99.0 %, Merck KGaA, Darmstadt, Germany) and Sodium Chloride (≥ 99.0, Merck KGaA, Darmstadt, Germany) were used to create solvents for producing fibrils and spherulites. All solvents were prepared with reverse-osmosis water (Milli-Q) as diluent and were filtered through a 0.1 µm PES filter mounted onto a 50 mL centrifuge tube (#514-0366, VWR, Radnor, PA, USA) before use. Zinc-complexed lyophilized human insulin was kindly provided by Novo Nordisk A/S. All pH adjustments were performed using 1 M HCl or 1 M NaOH (99.0 %, Merck KGaA, Darmstadt, Germany). All samples were filtered prior to stress application through a 0.1 µm inorganic membrane filter (#6809-2012, Whatman, Little Chalfont, UK) using silicone- and latex-free syringes (#4050-X00V0, Henke Sass Wolf, Tuttlingen, Germany) into 15 mL Nunc conical centrifuge tubes (#339651, Thermo Fisher Scientific, Waltham, MA, USA). Filters and syringes were pre-washed x3 with Milli-Q. A continuous flow of sample was ensured during filtration to avoid pre-aggregation [27].

### Identification of conditions for fibril and spherulite formation

Human insulin was dissolved in 20% acetic acid solution (v/v), containing either 0 mM or 500 mM NaCl, to a final protein concentration of 5 mg/mL. To reach the target value of pH 2.0, the preparations were adjusted with 1 M NaOH. Samples were filtered as described in the materials section. Aliquots of 1 mL sample were added to 1.5 mL Eppendorf tubes (#0030 120.086, Eppendorf AG, Hamburg, Germany) and placed into a thermomixer (thermomixer comfort, Eppendorf AG, Hamburg, Germany) at 60 °C, either quiescent conditions or with mechanical shaking 600 rpm for 24 hours.

### Preparation of amyloid-like fibrils

Human insulin was dissolved in 20% acetic acid solution (v/v) to a final protein concentration of either 0.5 or 5 mg/mL. To reach the target value of pH 2.0, the 0.5 mg/mL preparations were adjusted with 1 M NaOH. No pH adjustments were necessary for the 5 mg/mL samples. Sample preparations were filtered as described in the materials section. Aliquots of 1 mL sample were added to 1.5 mL Eppendorf tubes (#0030 120.086, Eppendorf AG, Hamburg, Germany) and placed into a thermomixer (thermomixer comfort, Eppendorf AG, Hamburg, Germany) at 60 °C, with mechanical shaking 600 rpm for 24 hours. For a detailed protocol, please refer to our previous work [27].

### Preparation of spherulites

Two different types of spherulites were prepared by adjustment of the solvent conditions under which they are formed. One type (AA spherulites) was prepared with 20% acetic acid (v/v), 300 mM NaCl and a final human insulin concentration of 1.75 mg/mL at pH 2.0. Another type (EtOH spherulites) was prepared with 40% ethanol (v/v), 250 mM NaCl with a final human insulin concentration of 5 mg/mL at pH 1.8. Both preparations were pH adjusted to reach the target pH values with 1 M NaOH. Samples were filtered as described in the materials section. Aliquots of 1 mL sample were added to 1.5 mL Eppendorf tubes (#0030 120.086, Eppendorf AG, Hamburg, Germany) and placed into a thermomixer (thermomixer comfort, Eppendorf AG, Hamburg, Germany) at 55 °C, quiescent conditions for 8 hours. For a detailed protocol, see our previous work [27].

### Flow Imaging Microscopy

Samples containing either amyloid-like fibrils or spherulites were analysed using a Micro-Flow Imaging microscope (model DPA 4200, BrightWell Technologies Inc., Kanata, ON, Canada). We introduced a method with no prior sample purging and a manual stop function. This was to ensure that the measurements could be instantly stopped without losing already analysed data e.g., in case of flow cell clogging (#4002-002-001, 100 µm, silane coating, ProteinSimple, Toronto, ON, Canada). Flow cell clogging can occur during analysis of spherulite samples, due to their tendency to interact with surfaces inside the flow cell. Prior to measurements, the flow cell was cleaned with 2% Hellmanex and Milli-Q until a baseline particle concentration of <150 #/mL could be detected. The illumination was automatically optimized by the instrument on a solvent equivalent to the one present in the analysed sample. AA spherulite samples were first homogenised by pipetting, then left for 10 minutes to allow the largest clusters to sediment. Supernatant containing individualized spherulites was used for analysis using MFI. EtOH spherulites and fibril samples were analysed without further sample preparation. Between measurements of different samples, the instrument was cleaned with 2% Hellmanex and Milli-Q. The flow cell was visually checked for protein particles clogged in the inlet. If clogging had occurred, the flow cell was removed from the instrument and manually cleaned with 2% Hellmanex and pressurized air. Data evaluation was performed using an in-house build software [28]. The first 250 frames were discarded from the analysis due to slight dilution of the samples in these frames. Images of the individual particles and their estimated Equivalent Circular Diameter (ECD, µm) were used for evaluating sample morphology, integrity, reproducibility, and homogeneity.

### Transmission Electron Microscopy

To increase surface hydrophilicity before sample addition, carbon filmed cobber 400 mesh grids (Ted Pella, Redding, CA, USA) were glow discharged for 30 seconds under vacuum using a Leica Coater ACE200 (Leica Microsystems, Wetzlar, Germany). For spherulite samples, the spherulite particles were left to sediment for 10 minutes into a pellet, then 3 µL of the supernatant was applied to the grids and left for 180 seconds. Fibril samples were deposited directly onto the grid to maintain sample integrity. However, the samples were only left on the grid for 3 seconds. Hereafter, the excess sample was removed by filter paper, directly followed by a 3x dropwise addition of 2% Uranyl Acetate (UA) solution (total volume 20 µL) for negative staining of the sample. Excess UA solution was removed in between drops and the grid was left to dry until analysis. A CM100 TWIN Transmission Electron Microscope (Phillips, Eindhoven, Netherlands) with tungsten emitter was used for all measurements. Image acquisition was performed via a Veleta camera (Olympus, Hamburg, Germany) using the ITEM software. The ImageJ software was used for length determination of the fibrils. Only fibrils, where start and end point could easily be detected in the TEM images, were included in the analysis. If the fibril length could not be appropriately described by a straight line, the fibril length was divided into smaller sections and subsequently summarized to obtain the full length. For each sample, 100 fibrils were measured to acquire adequate statistics.

### Aggregation kinetics

Kinetics of spherulite formation were investigated using a CLARIOstar plate reader (BMG Labtech, Ortenberg, Germany). Freshly prepared (unstressed) solutions for the formation of each spherulite type were mixed with Thioflavin T (∼65 %, T3516, Sigma Aldrich, Saint Louis, MO, USA) so that a final concentration of 20 µM of the dye was reached. Samples were transferred (200 µL/well) into 96-well plates black with clear-bottom (#265301, Thermo Fisher Scientific, Waltham, MA, USA). The samples were stressed at 55 °C and an excitation wavelength of 450 nm, emission wavelength 486 nm with a dichroic mirror at 465 nm were used to detect ThT fluorescence as a function of time. Moreover, a focal height of 2.6 and gain of 1000 were employed. Each well was subjected to 10 flashes pr. cycle with a cycle time of 309 seconds. The total number of cycles was set according to the duration of the aggregation reaction, which varied between the two spherulite types; minimum 10 hours for AA spherulites and minimum 24 hours for EtOH spherulites. The lag-time is defined as the EC10. It is determined as the time point, where 10% of the maximum fluorescence signal has been obtained.

### Confocal microscopy

Spherulite samples were either stained with Thioflavin T (∼65 %, T3516, Sigma Aldrich, Saint Louis, MO, USA) for amyloid content detection or dual-stained with 1,8 ANS (#A47, Molecular Probes Inc., Eugene, OR, USA) and SeTau-647-Maleimide (K9-4148, SETA BioMedicals, Urbana, IL, USA) for hydrophobicity/hydrophilicity detection, respectively. The staining was performed 24 hours before measurements. A microscope chamber slide was assembled by adhering a 6-channelled sticky slide VI 0.4 (#80608, Ibidi GmbH, Martinsried, Germany) onto a cover glass slide. To each channel, 80 µL stained sample was added. Imaging was performed on a confocal microscope (IX81, Olympus, Tokyo, Japan) using a 100x oil immersion objective (UPlanSApo 100x/1.4o oil, Olympus, Tokyo, Japan) and FV1000 camera (Olympus, Tokyo, Japan) at resolution 1024 × 1024 pixels. ThT fluorescence was acquired with the 458 nm laser, ANS fluorescence with the 405 nm laser and SeTau with the 635 nm laser. The ImageJ software was used for determination of spherulite size from the bright-field images. Since spherulites are not perfectly spherical, their characteristic size was defined as the longest axis. For each sample, 80 spherulites were measured to acquire adequate statistics.

### Environmental Scanning Electron Microscopy

Spherulite samples were stressed for 24 hours and centrifuged (5417R, Eppendorf, Hamburg, Germany) at 14,000 rpm, 4 °C for 30 minutes. The pellet from three tubes was pooled, to ensure sufficient material for analysis. A drop of spherulite sample was transferred onto a specimen stage and gently spread out to cover the surface. Measurements were performed with a FEI Quanta 3D FEG Scanning Electron Microscope (Thermo Fisher Scientific, Hillsboro, OR, USA) in environmental mode allowing imaging of hydrated samples with minimum sample preparation. A low-vacuum detector (Lv-SED) and acceleration voltage of either 5.00 kV or 10.00 kV were used for all acquisitions.

### Solvent exchange procedures

All samples were solvent exchanged. Two approaches for exchanging solvent were tested: dialysis and centrifugation. In the first approach, samples of fibrils and spherulites were dialysed for 48 hours at 4 °C against PBS using 100 kD MWCO Float-A-Lyzer G2 dialysis devices (#G235059, Spectra/Por, Spectrum Laboratories Inc., Rancho Dominguez, CA, USA). Prior to dialysis, the device membranes were pre-wetted to remove glycerine and optimize permeability according to manufacturer recommendations. First, soaking the membranes in 10% ethanol (v/v) for 10 minutes followed by soaking in Milli-Q for 30 minutes, under stirring. Then, 3 mL sample was added to each device and directly transferred to a beaker with 3 L PBS (phosphate-buffered saline, w/o calcium and magnesium, pH 7.4) under stirring. After 24 hours, the PBS was discarded and replenished with fresh PBS. Dialysis proceeded under stirring for 24 hours, before sample withdrawal. In the centrifugation approach, samples of fibrils were centrifuged (5417R, Eppendorf, Hamburg, Germany) at 14,000 rpm, 4 °C for 3 hours. The supernatant was discarded, and PBS was added. The pellet containing the aggregates was gently resuspended by pipetting in PBS. The procedure was repeated x3, with centrifugation time of 1 hour, to ensure complete exchange of solvent.

### Fourier-Transform InfraRed Microscopy

Spherulites, in either original solvent or PBS, were immobilized between two CaF2 windows (13 mm diameter, 0.5 mm thickness, Crystran Ltd., Dorset, UK) and placed into a custom-made compression cell utilizing a similar procedure as described by Schack *et al*. [29]. With this setup, the spacing between the two CaF2 windows was expected to be approximately 4.5 µm. Measurements were performed on a Nicolet iN10 MX FTIR microscope (Thermo Fisher Scientific, Waltham, MA, USA) equipped with a single-element MCT-A detector, cooled with liquid nitrogen. For each sample preparation, ten individualized particles, which entirely covered an aperture size of 25 µm x 25 µm, were selected. Prior to measurements, a background spectrum of bulk solution with similar aperture size was acquired and automatically subtracted from the particle spectrum by the OMNIC Picta software (Thermo Fisher Scientific, Waltham, MA, USA). FTIR spectra were collected in transmission mode in the range 4000 cm-1 to 675 cm-1. Each spectrum is an average of 512 scans with spectral resolution of 4 cm-1. The interferograms were processed with a Blackman-Harris 3-terms apodization function with a zero filling factor of one. The amide I peak (1600 cm-1 to 1700 cm-1) was used to evaluate secondary structural differences between the spherulite samples.

## Results and Discussion

### Salt content and mechanical stress affect fibril and spherulite formation

Fibrils and spherulites are both commonly formed under highly acidic conditions at elevated temperatures, and they can co-exist [17, 18]. This makes it difficult to prepare samples containing only fibrils or spherulites. We investigated the effect of salt concentration (0 mM NaCl or 500 mM NaCl) and shaking speed (0 rpm or 600 rpm), which are parameters known to affect fibril or spherulite formation [4, 17, 20, 30]. Here, the parameters were investigated in a formulation consisting of 5 mg/mL human insulin in 20% acetic acid solution at pH 2.0 and the samples were stressed at 60 °C for 24 hours. The presence of fibrils was evaluated by TEM, while spherulites were evaluated by MFI.

The results demonstrate the importance of salt in the formulation to favour spherulite formation rather than fibril growth (Fig. 1). The fibrils appear as elongated thin fibres on the TEM grid, while the spherulites appear as dark clusters of spherical aggregates in the MFI. In fact, the presence of salt led to a drastic decrease in fibril propensity. We argue that the increase in ionic strength, by addition of sodium chloride, shields the repulsive charges between insulin molecules favouring spherulite formation. Here initial coalescence processes are likely dominating, leading to formation of a core from where fibrils can radially grow. In the absence of salt, the repulsion between the highly charged insulin molecules is strong. Here the aggregation process is mainly driven by an initial conformational instability, from which fibrils form [31]. The observed effect has also been reported by Waugh et al. [32] for bovine insulin in phosphoric acid solutions with or without salt. Nevertheless, the effect of salt does not seem to be apparent for neither bovine insulin nor human insulin in aqueous solutions with hydrochloric acid. Under these conditions, spherulites can readily form both with and without salt present, where they co-exist with free fibrils [4, 11]. We speculate whether the difference originates from the initial oligomerization state of the protein, affecting the aggregation process. In acetic acid, insulin primarily exists in its monomeric form [33], whereas in hydrochloric acid it primarily exists in a monomer/dimer equilibrium [34, 35].

**Fig. 1.**
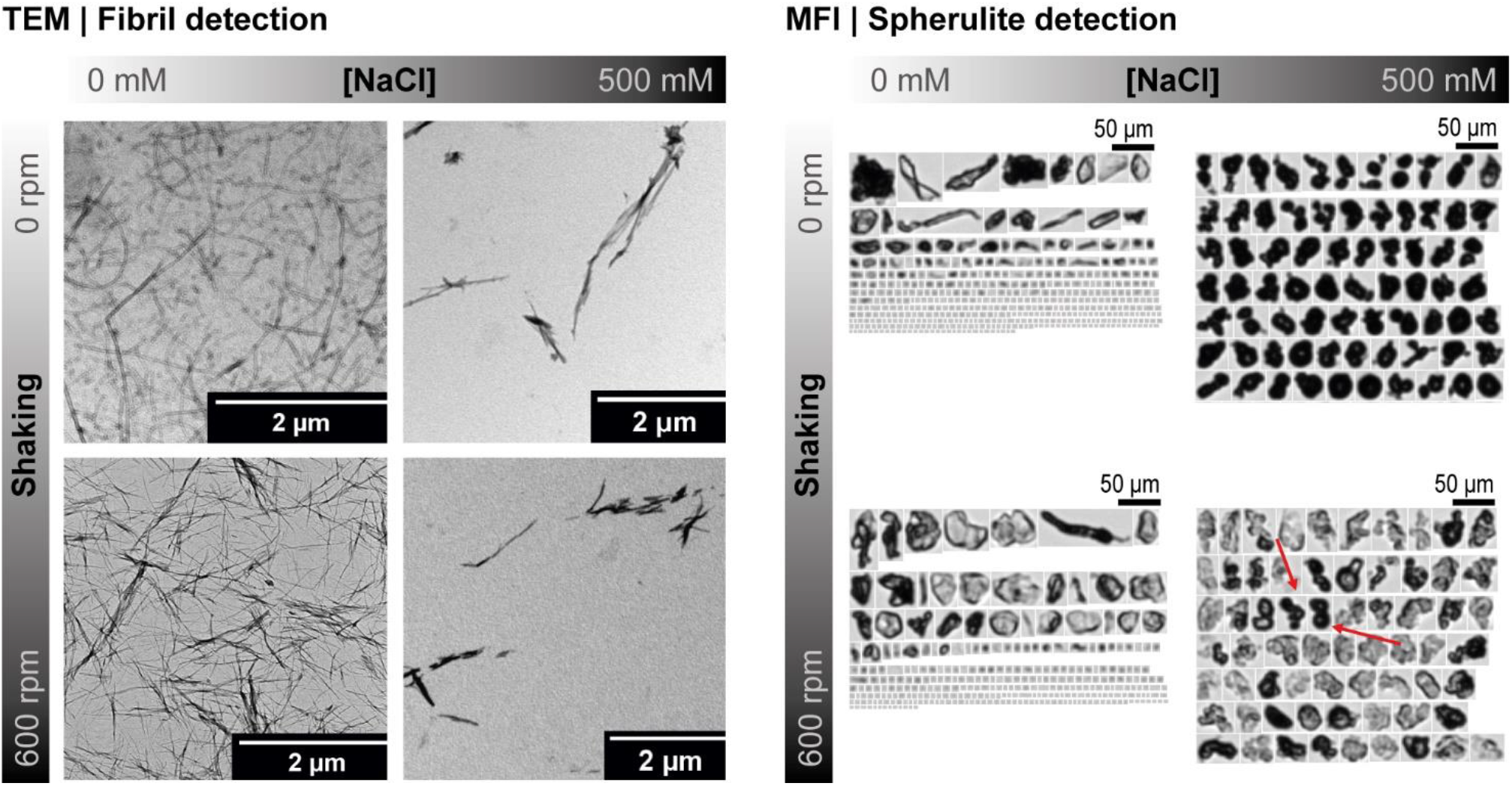
Effect of NaCl concentration and shaking speed on the formed aggregate types. Samples containing 5 mg/mL human insulin in 20% acetic acid at pH 2, heat stressed at 60 °C for 24 hours, either with 0 mM NaCl or 500 mM NaCl present in the formulation and either at quiescent conditions (0 rpm) or mechanical shaking (600 rpm). Representative images of the samples analysed with Transmission Electron Microscopy (left panel) for detection of fibrils and Micro-Flow Imaging (right panel) for detection of spherulites. In the MFI panel, all particles >1 µm are displayed for samples without NaCl, while particles in the size range 24-30 µm are displayed for samples containing 500 mM NaCl to illustrate the representative morphology. Red arrows indicate the presence of a few spherulites formed in the specific sample.

Moreover, we observed that shaking supress spherulite formation and rather promote production of amorphous aggregates. This is consistent with recent results by Pounot *et al*. [17] for human insulin in aqueous solutions with hydrochloric acid, where it was argued that the mechanical perturbation prevents formation of sufficiently stable nuclei during spherulite development. Fibrils formed both under shaking and quiescent conditions, where shaking produced shorter fibrils compared to fibrils formed at quiescent conditions (Fig. 1). Additionally, we found that fibrils formed during shaking were in general more reproducible in terms of length and morphology, compared to fibrils formed at quiescent conditions. The effect of shaking on fibril length has previously been observed by Yoshihara *et al*. [13] for human insulin and Arosio *et al*. [36] for β-LAC protein. Yet, the aggregation mechanism controlling fibril growth when mechanical stress is applied and the effect on the resulting length are contradictory for the two publications, illustrating the complexity of the process. We did not further investigate the mechanism of fibril growth as it falls out of the scope of this work.

Our data showed that by controlling salt concentration and mechanical stress, we were able to form samples containing a single aggregate type. A sample consisting of spherulites can be obtained from a 20% acetic acid formulation containing 500 mM NaCl, which is heat stressed (60 °C) at quiescent conditions. While a sample consisting of fibrils can be obtained by absence of NaCl. Long or short fibrils can be formed under quiescent conditions or under mechanical shaking at 600 rpm, respectively.

### Initial protein concentration impacts fibril length

Having identified a condition where only fibrils are formed, we wanted to create samples which would allow for investigation of the effect of specific morphological characteristics in biological systems such as e.g., immunogenicity assessment. Aggregates have been suggested to bind receptors on antigen presenting cells, which recognizes repetitive epitopes and hereby initiate an immune response through pathogen-associated molecular patterns (PAMPs) [37]. From a cellular perspective, fibrils could be viewed as repetitive epitopes, thus the length of the fibrils could potentially influence the immunogenicity.

Van Raaij et al. [38] suggested that since fibril formation is considered to be controlled by a nucleation-polymerization process, the initial protein concentration would be expected to affect the length of α-synuclein fibrils. Their results showed that both fibril length and the number of fibrils were highly concentration dependent. A low initial protein concentration could result in fewer stable nucleation points for the fibrils to grow from, leading to less fibrils being formed. Additionally, less protein will be available in solution in the growth-phase, leading to shorter fibrils.

If the observations about protein concentration and fibril length from Van Raaji et al. [38] translates to human insulin in acetic acid, we could potentially use this to control fibril length. In the further investigation of fibril length, the most reproducible procedure, namely: 5 mg/mL human insulin in 20% acetic acid at pH 2.0, stressed by exerting heat (60 °C) and mechanical shaking (600 rpm) for 24 hours was applied. Protein concentration was decreased 10-fold from 5 mg/mL to 0.5 mg/mL and the fibril morphology and length were evaluated by TEM.

Our results showed, that decreasing the protein concentration 10-fold resulted in shorter fibrils with a mean length of 300 nm, as compared to 1.2 µm for the high protein concentration sample, with otherwise similar appearance (Fig. 2). The heterogenous dispersion of fibrils on the TEM grid did not allow for a quantitative evaluation of the number of fibrils formed for each concentration. However, it was qualitatively evaluated by visual inspection of the samples after stress. Here it was observed that less fibril material (appearing like an opalescent gel-phase) had been formed in the 0.5 mg/mL sample compared to the 5 mg/mL sample, as expected. Both samples were analysed for potential co-existence of spherulites (Fig. S1). Only particles attributed to material impurities were found, indicating that the samples indeed are composed of only fibrils.

**Fig. 2.**
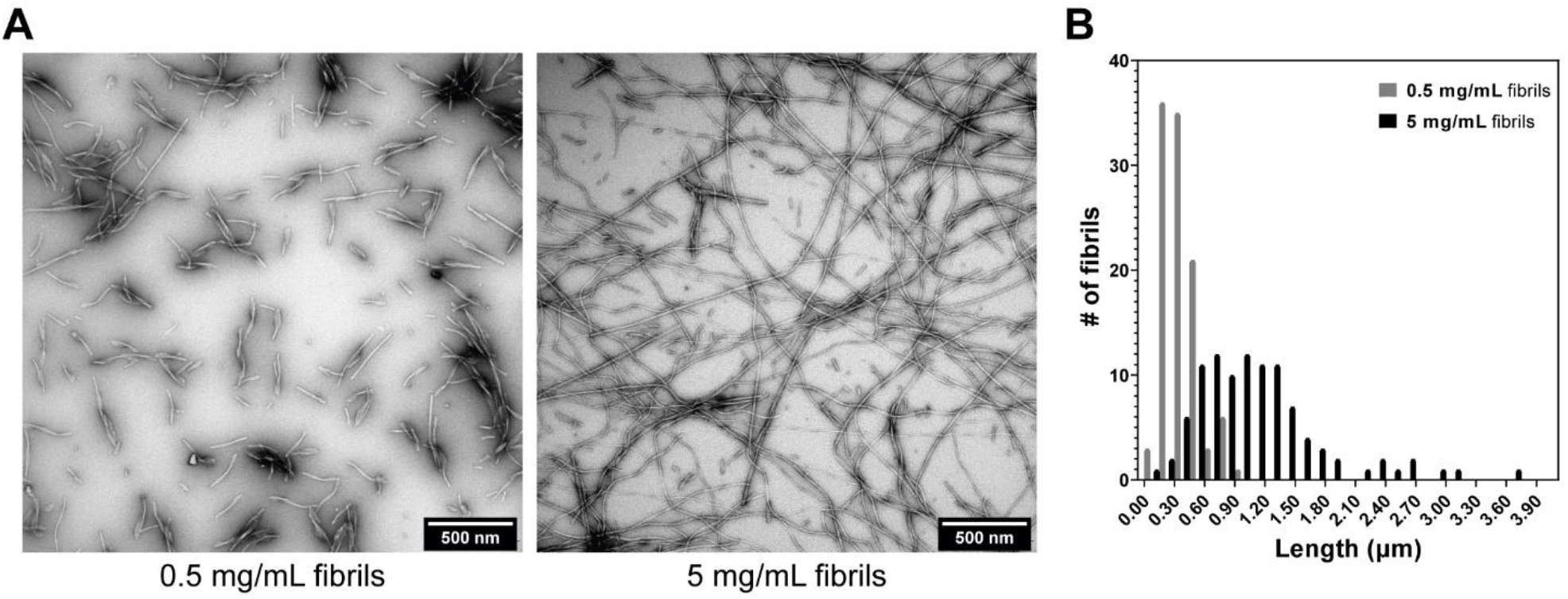
Protein concentration and its effect on fibril length. A) Representative Transmission Electron Microscopy (TEM) images of fibrils formed by either 0.5 mg/mL (left) or 5 mg/mL (right) human insulin in 20% acetic acid, pH 2.0 at 60 °C and 600 rpm for 24 hours. B) The difference in fibril length is quantified by ImageJ analysis from N = 100 fibrils for each protein concentration.

### Solvent polarity controls spherulite secondary structure

Having identified a condition where only spherulites are formed, we wanted to create samples which would allow for investigation of the effect of specific morphological characteristics in biological systems such as e.g., immunogenicity assessment. Aggregates consisting of misfolded proteins and amyloid-like structure may result in the break of immune tolerance in transgenic mice [39]. Amyloid-like spherulites produced in acetic acid contain a high amount of aggregated β-sheet, compared to native human insulin [11, 14], thus reducing the amount of aggregated β-sheet present in the spherulite could potentially reduce the immunogenicity risk of this aggregate type.

*Vetri et al*. [40] showed that spherulites with a more native-like secondary structure (decreased content of aggregated β-sheets) can be formed in the presence of 40% ethanol, 0.25 M NaCl at pH 1.8 compared to 0% ethanol. Spherulites were formed under similar conditions in our lab and the particle characteristics was compared to spherulites formed in acetic acid using an optimized protocol, which could reproducibly form individualized spherulites [27]. The final formulation contains 1.75 mg/mL human insulin in 20% acetic acid with 0.3 M NaCl adjusted to pH 2.0. The stress temperature was decreased from 60 °C to 55 °C and the stress time from 24 hours to 8 hours since we observed particle-particle clustering after this time point. For further reference, the spherulites formed in acetic acid are called AA spherulites, while spherulites formed in ethanol are called EtOH spherulites.

Ethanol preferentially binds to the protein, modifies the surrounding hydration layer, and increases the repulsion between molecules. This results in a delay of the aggregation process, which is observed as a prolonged lag-phase in the kinetics [40, 41]. A similar effect was observed, when comparing the kinetics for AA spherulites and EtOH spherulites (Fig. S2). Here the lag-phase was increased from 2.4 hours to 10.6 hours, respectively. We examined the supernatant of the two samples utilizing TEM, to confirm that the decreased salt concentration in the spherulite samples, compared to our initial investigation, would not increase the propensity of fibrils. As expected, only a few fibrils were observed on the grid indicating that the samples indeed contain primarily a single species of spherulites (Fig. S3).

Next, we characterized the morphology of the two spherulite types. Here representative spherulites from both samples show a spherical appearance with bright-field microscopy and SEM (Fig. 3 and Fig. 4A). The Maltese cross pattern displayed under cross-polarized light indicates that the particles contain specific internal geometrical arrangement and the positive fluorescence response upon staining with Thioflavin T suggests the presence of amyloid-like structure (Fig. 3); all classic morphological characteristics of spherulites. Moreover, the average particle diameter for the AA spherulites and EtOH spherulites were determined to be 24 µm and 17 µm, respectively, demonstrating a size similarity between the two samples (Fig. 4B). Interestingly, upon dual staining of the particles with a hydrophilic (SeTau) and hydrophobic (ANS) dye, the AA spherulites present a hydrophilic environment throughout the entire particle with a hydrophobic rim, whereas the EtOH spherulites present a hydrophobic inner environment with a hydrophilic rim. The differences in the hydrophilicity/hydrophobicity seem to be linked to a more open spherulites structure for the AA Spherulites with a hydrophilic environment throughout the particle and a more hydrophobic core for the more compact EtOH Spherulites. *Vetri et al*. found the hydrophilicity to be distributed uniformly throughout the EtOH spherulites and not only residing in the rim. The discrepancy could be due to the use of a different hydrophilic dyes, Alexa-647 vs. SeTau-647, or that dual staining with ANS instead of ThT (used by Vetri et al.) could induce a different competition between dye molecules leading to a different staining pattern.

**Fig. 3.**
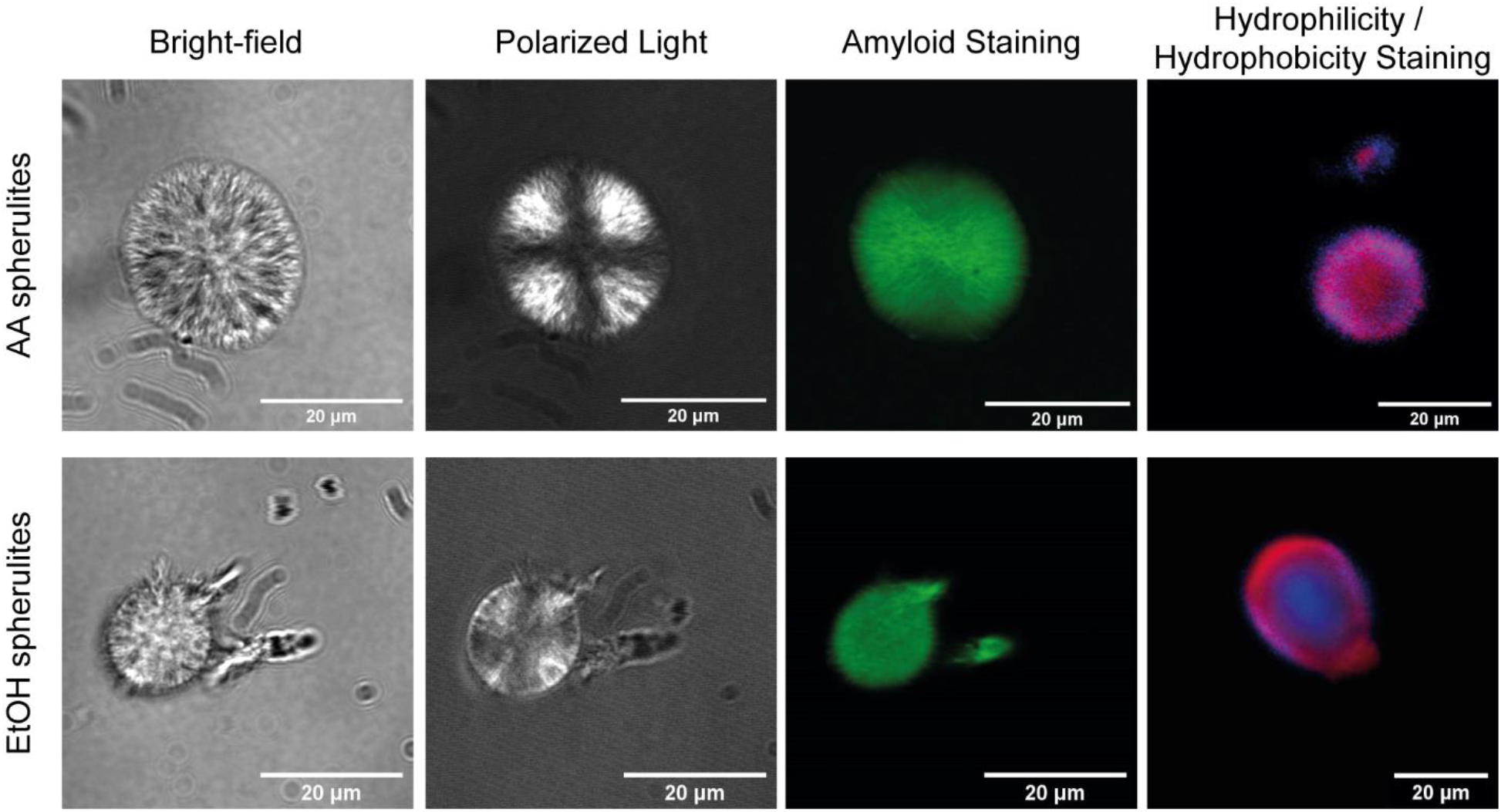
Morphological analysis of spherulites with confocal microscopy. Spherulites are formed in either 20% acetic acid, 0.3 M NaCl pH 2.0 (top) or 40% ethanol, 0.25 M NaCl pH 1.8 (bottom) by stress at 55 °C, quiescent conditions for 8 hours. The morphological features of the two spherulite types were investigated by bright-field microscopy (left), polarized light (middle left), staining with amyloid sensitive dye Thioflavin T (middle right) and two-color staining (right) with ANS (blue) and SeTau (red). The two-color staining was used to indicate hydrophobic (ANS) and hydrophilic (SeTau) areas on the spherulites. See Fig. S4, where the two-color staining has been split into two channels instead of overlay

**Fig. 4.**
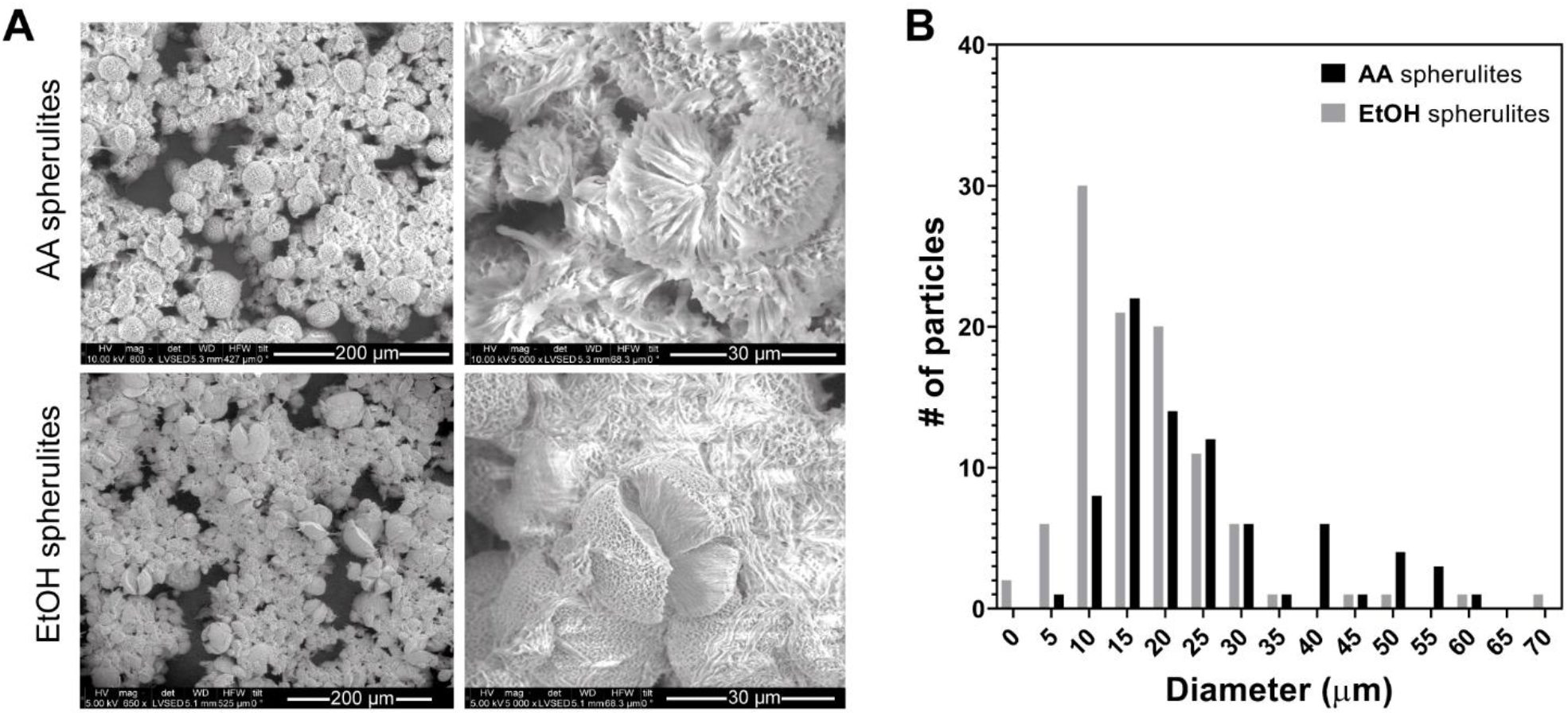
Morphological analysis with Scanning Electron Microscopy and size determination of spherulites. A) Morphological analysis with SEM on AA spherulites formed in 20% acetic acid, 0.3 M NaCl pH 2.0 (top) or EtOH spherulites formed 40% ethanol, 0.25 M NaCl pH 1.8 (bottom) formed at 55 °C, quiescent conditions for 24 hours. A zoom on a single spherulite from each population (right) shows fibrils radially forming from an inner nucleus. The scalebar for the overview images (left) is 200 µm and the scalebar for the zoom-in images (right) is 30 µm. B) Size determination of spherulites from bright-field confocal images of the two spherulite types estimated by ImageJ analysis. Distributions comprise 80 individualized spherulites for each type. AA spherulites are black bars, whereas EtOH spherulites are grey bars.

Lastly, we examined the secondary structure of single spherulites from each sample by FTIR microscopy (Fig. 5A). Surprisingly, the FTIR spectra for the EtOH spherulites exhibited an increased aggregated β-sheet content compared to the AA spherulites, in contradiction to the results reported by Vetri *et al*. We speculate whether the discrepancy in the results could be due to a solvent exchange step from the original formulation to D2O just prior to their FTIR measurements. This hypothesis is further elaborated in the next section.

**Fig. 5.**
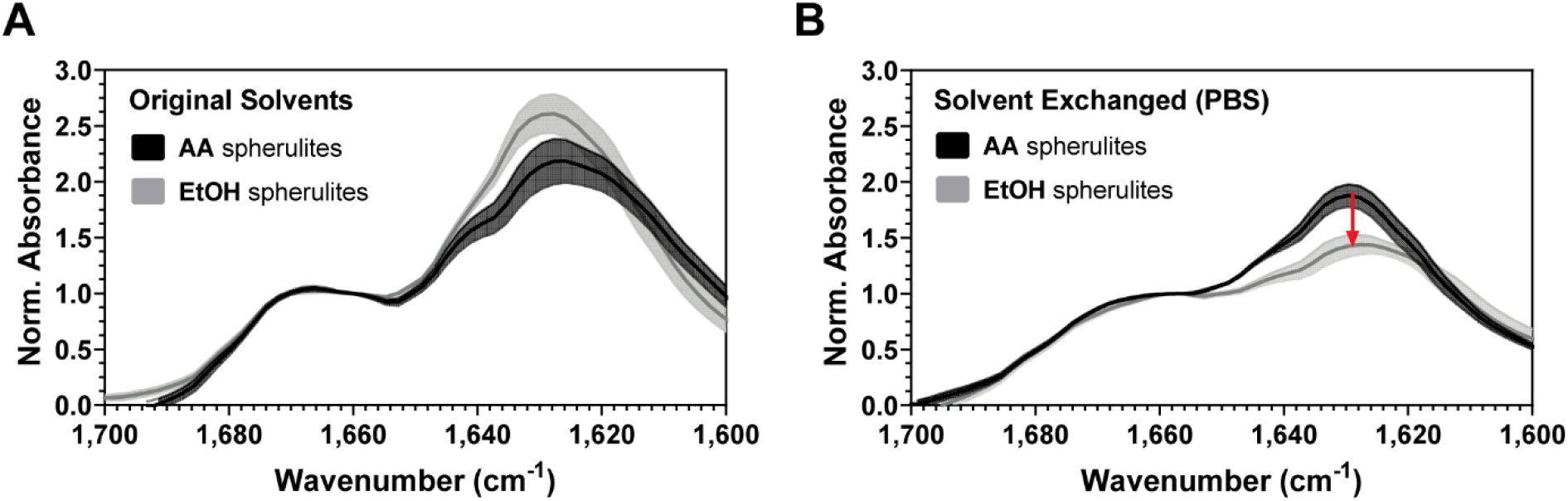
Fourier-Transform InfraRed Microscopy measurements on single spherulites. The FTIR spectrum of either AA spherulites (black) or EtOH spherulites (grey) were obtained in either A) the original solvent (20% acetic acid, 0.3 M NaCl, pH 2.0 or 40% ethanol, 0.25 M NaCl, pH 1.8) or B) after dialysis in PBS (w/o calcium and magnesium, pH 7.2-7.4). The shaded error bars denote standard deviation from ten individual particles (25×25 µm2). The spectra are normalized to 1656 cm-1, where α-helix content resides, to emphasize changes in aggregated β-sheet content observed at 1627 cm-1. The red arrow indicates a drastic decrease in aggregated β-sheet content for EtOH spherulites after solvent exchange to PBS.

### Solvent exchange affects morphological integrity

When performing *in vitro* aggregation assays, the formulation conditions are often incompatible with most biological assays. Solvent exchange is often readily applied without further notice on the impact it could have on the aggregates. Yet, exchanging the surrounding solution could potentially affect the morphological integrity of the aggregates due to the drastic change in formulation composition. To investigate the effect of solvent exchange on the morphological integrity and secondary structure of our produced fibril and spherulite samples, the sample solvents were exchanged to phosphate buffer saline (PBS, pH 7.2-7.4), a commonly used buffer in e.g., cell experiments. We tested two commonly used solvent exchange methods: centrifugation and dialysis. The fibril integrity was evaluated by TEM, while spherulite integrity was evaluated by MFI.

The results showed that fibril morphology was completely disrupted for the 0.5 mg/mL fibril sample utilizing any of the tested solvent exchange methods (Fig. 6). For the 5 mg/mL fibril sample, the length was clearly truncated. Additionally, massive clustering effects were observed, especially for the centrifugation method. The secondary structure of the fibrils after solvent exchange was not investigated due to the poor morphological resemblance to the original samples. The fibril samples were deemed unsuitable for further use in biological assays. The centrifugation method was not used for the spherulites since it was not possible to reliably pellet EtOH spherulites without substantial particle loss.

**Fig. 6.**
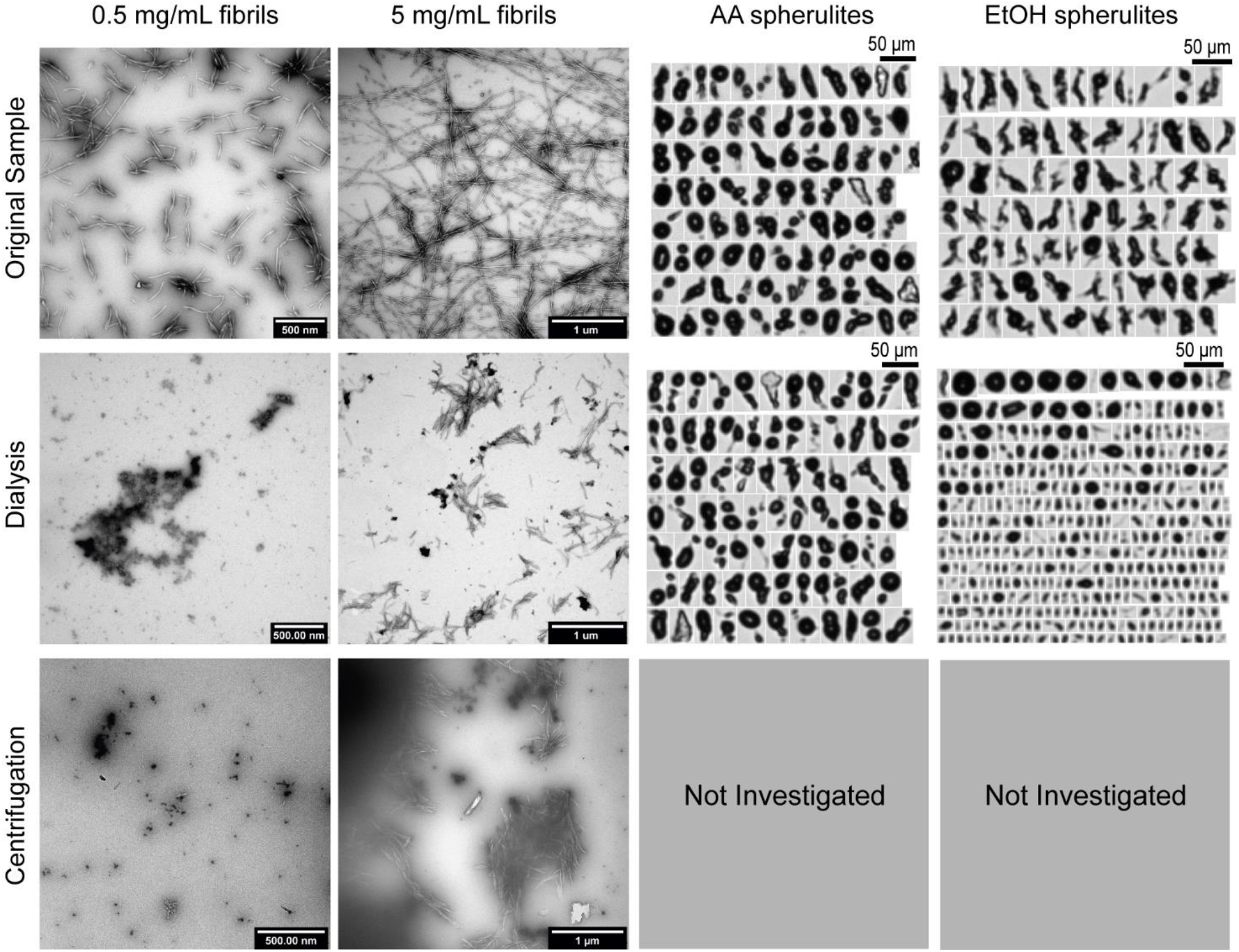
Different solvent exchange methods and their impact on morphological integrity. Representative images from samples in either its original solvent (acetic acid or ethanol) or solvent exchanged samples (PBS) are shown for direct comparison. The fibril integrity is evaluated by Transmission Electron Microscopy, whereas spherulite integrity is evaluated by Micro-Flow imaging.

Regardless, the spherulite morphology proved to be unaffected by the dialysis method. Thus, we recommend dialysis as a gentle method to solvent exchange spherulite samples for biological assays. FTIR microscopy on single EtOH spherulites after solvent exchange showed a drastically decreased content of aggregated β-sheet compared to the sample in original solvent (Fig. 5B). Whereas the AA spherulite secondary structure remained overall unaffected. These results are in line with Vetri *et al*. and supports our hypothesis that the solvent exchange step indeed affects the EtOH spherulite secondary structure.

## Conclusion

In this study, we reported that the balance between fibril and spherulite formation can be controlled for human insulin in acetic acid by the presence of salt. Samples consisting only of fibrils can be produced in absence of salt (NaCl), while salt is crucial for producing samples containing only spherulites. Moreover, fibril length can be tuned by either utilizing mechanical shaking during stress application or decreasing the initial protein concentration. Spherulites formed in ethanol contains more β-sheet structure, than spherulites formed in acetic acid. However, upon solvent exchange the β-sheet content decreased significantly in ethanol spherulites, whereas acetic acid spherulites remained unaffected. Also, fibril morphology displayed dramatic changes upon solvent exchange.

The profound impact of solvent exchange on the resulting morphology of aggregates has to our knowledge not been previously reported, yet its implications potentially affect conclusions drawn for aggregate characteristics and their corresponding biological outcome. We therefore strongly encourage characterization of samples both before and after solvent exchange, especially upon correlation between specific morphological characteristics of protein aggregates and biological outcomes, such as immunogenicity risk assessment. In a more general perspective, our findings reveal a key, hitherto unknown, element of the effect of the sample preparation and the post-production treatments, that may represent a source of artefacts. This fact is intimately related to the stability of the aggregates, a feature often overlooked in current studies and can potentially lead to misconception in the field.

## Acknowledgements

CT, MBN and MG acknowledge Novo Nordisk A/S for project funding and providing protein material. VF, AMS and CT acknowledge VILLUM FONDEN for supporting the project via the Villum Young Investigator Grant “Protein Superstructures as Smart Biomaterials (ProSmart)” 2018−2023 (Grant 19175). NSH and CT acknowledge VILLUM FONDEN (Grant 18333). The authors thank the VILLUM FONDEN (Grant 19175) for funding the CLARIOstar plate reader. For use of TEM and SEM, authors acknowledge the Core Facility for Integrated Microscopy, Faculty of Health and Medical Sciences, University of Copenhagen and Jesper Søndergaard Marino from Novo Nordisk A/S for use of MFI image evaluation software.

