## Supplementary information for "Morphological integrity of insulin amyloid-like aggregates depends on preparation methods and post-production treatments"

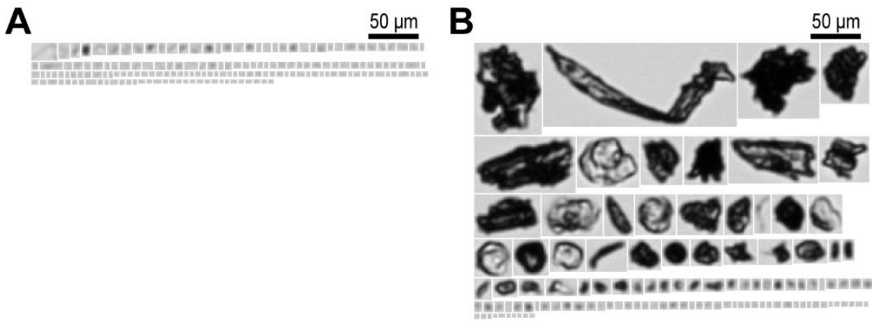


**Fig. S1 Micro-Flow Imaging analysis of fibril samples showing no spherulites have formed**. All particles >1 µm is shown. Samples containing either A) 0.5 mg/mL human insulin or B) 5 mg/mL human insulin in 20% acetic acid, pH 2.0, stressed at 60 °C, 600 rpm shaking speed for 24 hours.

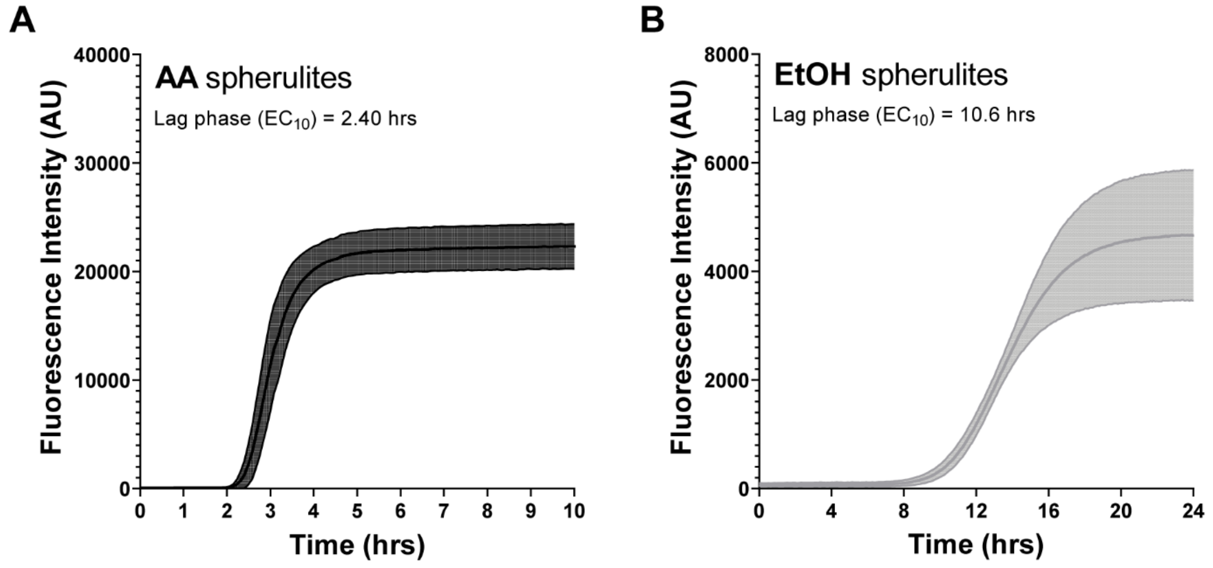


**Fig. S2 Kinetics of spherulite formation.** Human insulin in either A) 20% acetic acid, 0.3M NaCl pH 2.0 or B) 40% ethanol, 0.25M NaCl pH 1.8. Fluorescence intensity of the amyloid sensitive dye Thioflavin T was followed as a function of time as reporter of spherulite production. The initial lag-phase is defined by the $EC_{10}$ to 2.4 hours for AA spherulites and 10.6 hours for EtOH spherulites. The fluorescence intensity at t=0 was used to normalize the spectra. The shaded error bars are standard deviation of five independent measurements.

AA spherulite supernatant

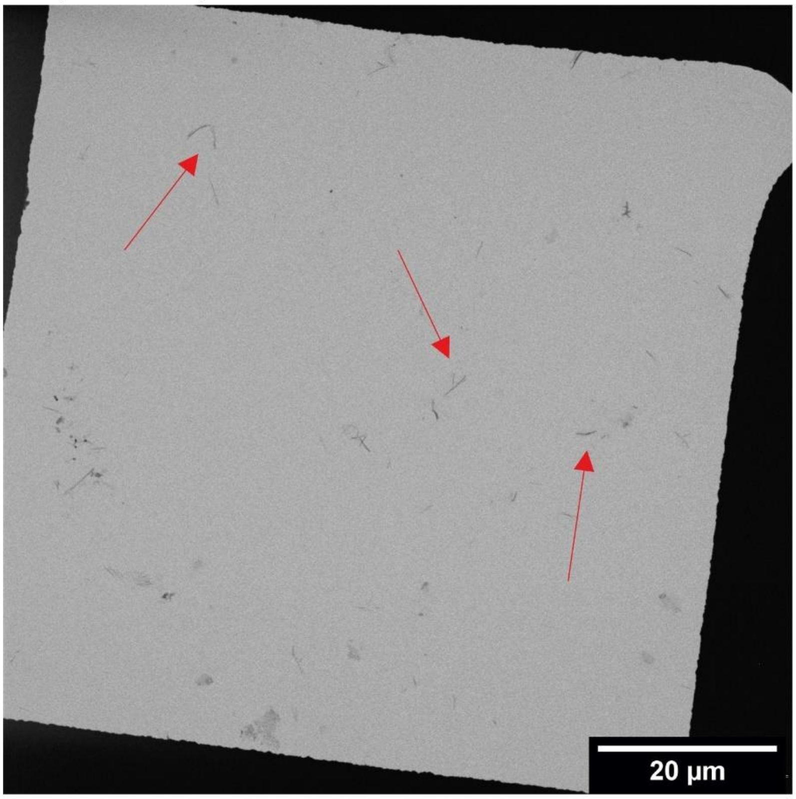


EtOH spherulite supernatant

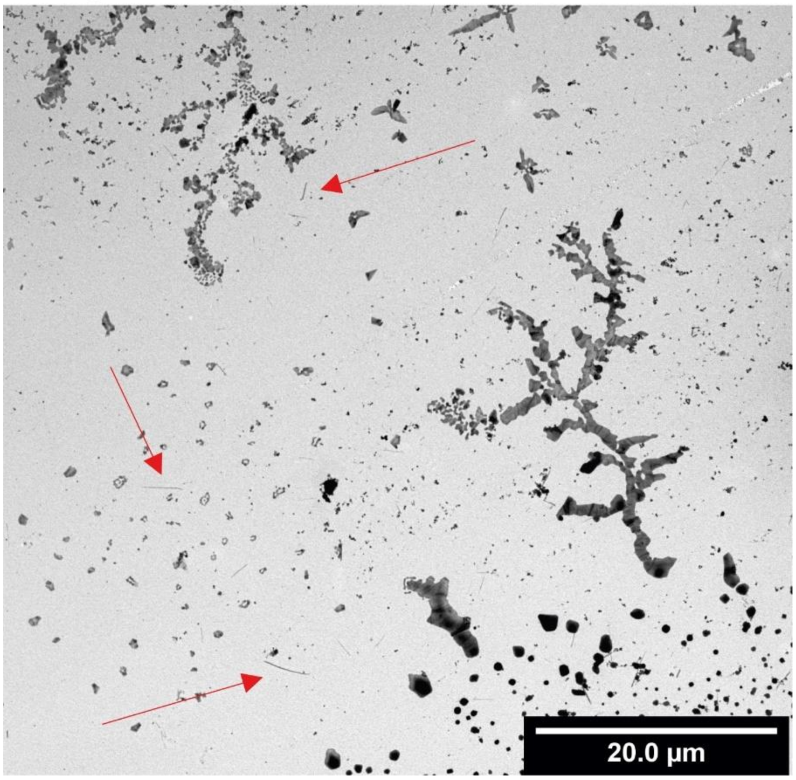


**Fig. S3 Representative Transmission Electron Microscopy images of the spherulite sample supernatants.** AA spherulites (top) and EtOH spherulites (bottom) generated by applied heat stress at 55 °C, quiescent conditions for 8 hours. Red arrows indicate the few fibrils found in either of the samples.

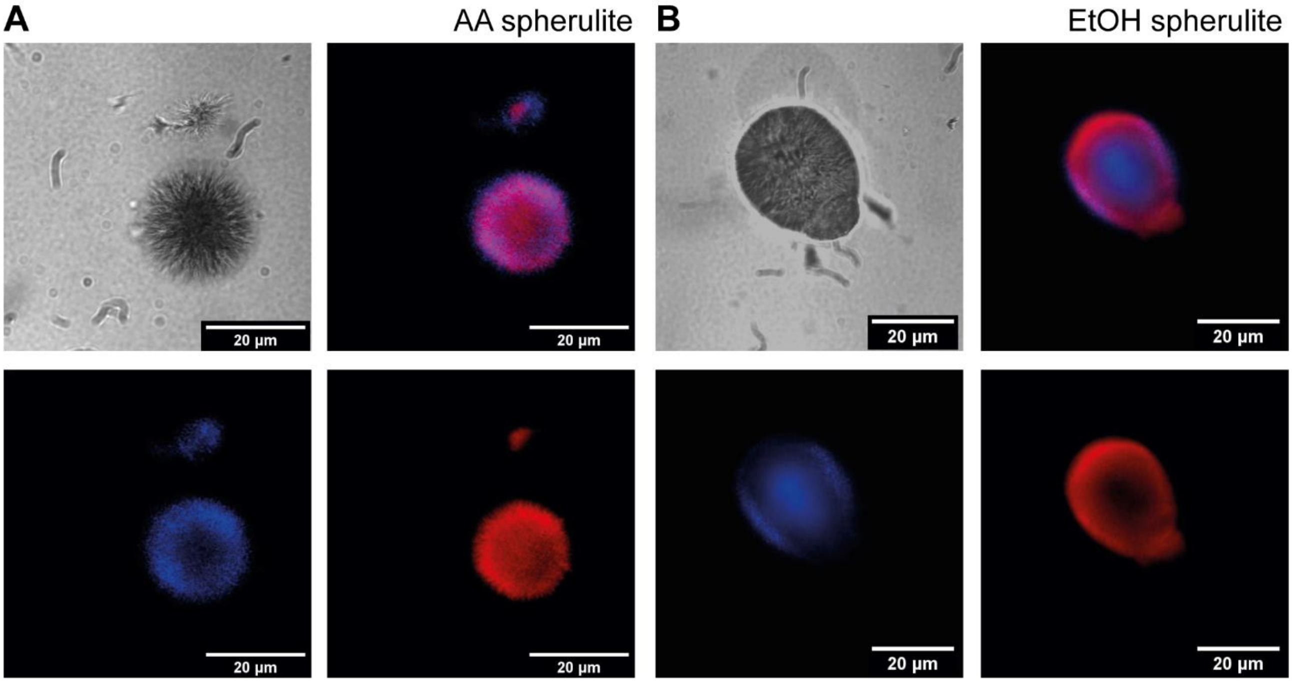


**Fig. S4 Dual staining of spherulites with confocal microscopy.** Spherulites are formed in either A) 20% acetic acid, 0.3 M NaCl pH 2.0 or B) 40% ethanol, 0.25 M NaCl pH 1.8 by stress at 55 °C, quiescent conditions for 8 hours. Bright-field images (top left), dual-staining overlay (top right), hydrophobicity staining with ANS (bottom left) and hydrophilicity staining with SeTau (bottom right). ANS signal is coloured blue, while SeTau signal is coloured red.
